# The unreasonable effectiveness of equilibrium gene regulation through the cell cycle

**DOI:** 10.1101/2023.03.31.535089

**Authors:** Jose M. G. Vilar, Leonor Saiz

## Abstract

Systems like the prototypical *lac* operon can reliably hold the repression of transcription upon DNA replication across cell cycles with just ten repressor molecules per cell and, in addition, behave as if they were at equilibrium. The origin of this type of phenomena is still an unresolved question of major implications. Here, we develop a general theory to analyze strong perturbations in quasi-equilibrium systems and use it to quantify the effects of DNA replication in gene regulation. We find a scaling law that connects actual transcription with its predicted equilibrium values in terms of a single kinetic parameter. We show that even the simplest, exceptionally reliable natural system functions beyond the physical limits of naïve regulation through compensatory mechanisms that suppress nonequilibrium effects. We validate the approach with both *in vivo* cell-population and single-cell characterization of the *lac* operon. Analyses of synthetic systems without adjuvant activators, such as the cAMP receptor protein (CRP), do not show this reliability. Our results provide a rationale for the function of CRP, beyond just being a tunable activator, as a mitigator of cell cycle perturbations.

## Introduction

Gene regulation depends to a large extent on equilibrium binding interactions (Ptashne and Gann, 2002). As such, equilibrium approaches, although do not consider explicitly many essential processes, have performed exceedingly well in both qualitative and even quantitatively understanding the major features of gene expression. They have been successfully widespread in modeling gene regulation since early pioneering work (Ackers et al., 1982; von Hippel et al., 1974) and have been applied to both prokaryotes (Guharajan et al., 2021; Kinney et al., 2010; Kuhlman et al., 2007; Lagator et al., 2022; Vilar and Leibler, 2003; Vilar and Saiz, 2005) and eukaryotes (Bashor et al., 2019; Carey et al., 1990; Gertz et al., 2009; Ptashne and Gann, 2002; Vilar and Saiz, 2011). The underlying assumption is that the equilibrium binding patterns, such as those observed in biochemical *in vitro* experiments, can be extrapolated to a living growing and multiplying cell. These patterns determine the transcriptional state of the system and dictate the amplitude of the outcomes of downstream nonequilibrium processes.

There are multiple cases in which it is required to move towards nonequilibrium kinetic approaches to obtain sensible results (Coulon et al., 2013; Doudna et al., 2017; Wong and Gunawardena, 2020). These include, for instance, cyclical processes based on both activating and repressive epigenetic mechanisms (Metivier et al., 2003), alternating transcription factor (TF) phosphorylation patterns (Kim and O’Shea, 2008), or even the long-term coexistence of induction states in the *lac* operon (Novick and Weiner, 1957; Ozbudak et al., 2004; Vilar et al., 2003). But, on the other hand, there are many complex situations with underlying energy expenditure, such as the patterning of drosophila embryos (Sayal et al., 2016; Segal et al., 2008), the cooperative binding of TFs as a result of nucleosome displacement (Mirny, 2010), and transcription regulation by biomolecular condensates (Hnisz et al., 2017), that have successfully been tackled with equilibrium-based approaches.

The effectiveness of equilibrium approaches has been quantified extensively in the *E. coli lac* operon and its synthetic derivatives. They have been tested across cell-population experiments for different strains, mutations, and synthetic constructs over a dynamic range in transcription rate changes larger than 10,000 (Garcia and Phillips, 2011; Kuhlman et al., 2007; Oehler et al., 2006; Saiz and Vilar, 2008; Vilar, 2010; Vilar and Leibler, 2003). Over this whole range, the typical discrepancies from the equilibrium predictions are, on average, just a factor 1.7 above or below the actual values (Vilar and Saiz, 2013a). Essentially this magnitude, a factor of 1.4, has also been observed in direct single-cell measurements in fast-growing microcolonies, which has been reasoned to be likely due to nonequilibrium effects in transcription initiation (Hammar et al., 2014).

In all cases, an unavoidable major far-from-equilibrium event is DNA replication, in which the promoter and DNA regulatory elements disappear momentarily upon the passage of the replication fork and reappear in duplicate (Fraser and Nurse, 1978; Voichek et al., 2016). During this time interval, there are no regulatory sequences to bind nor promoter to transcribe. After the promoter reconstitution, TFs and other regulatory molecules should find their duplicated binding sites fast enough for the equilibrium approach to hold (Fig. 1). Experimental evidence is being accumulated that indicates that the underlying processes are not sufficiently fast, even in the simplest systems (Elf et al., 2007; Hammar et al., 2014; Larson et al., 2011; Wheat et al., 2020).

**Figure 1.**
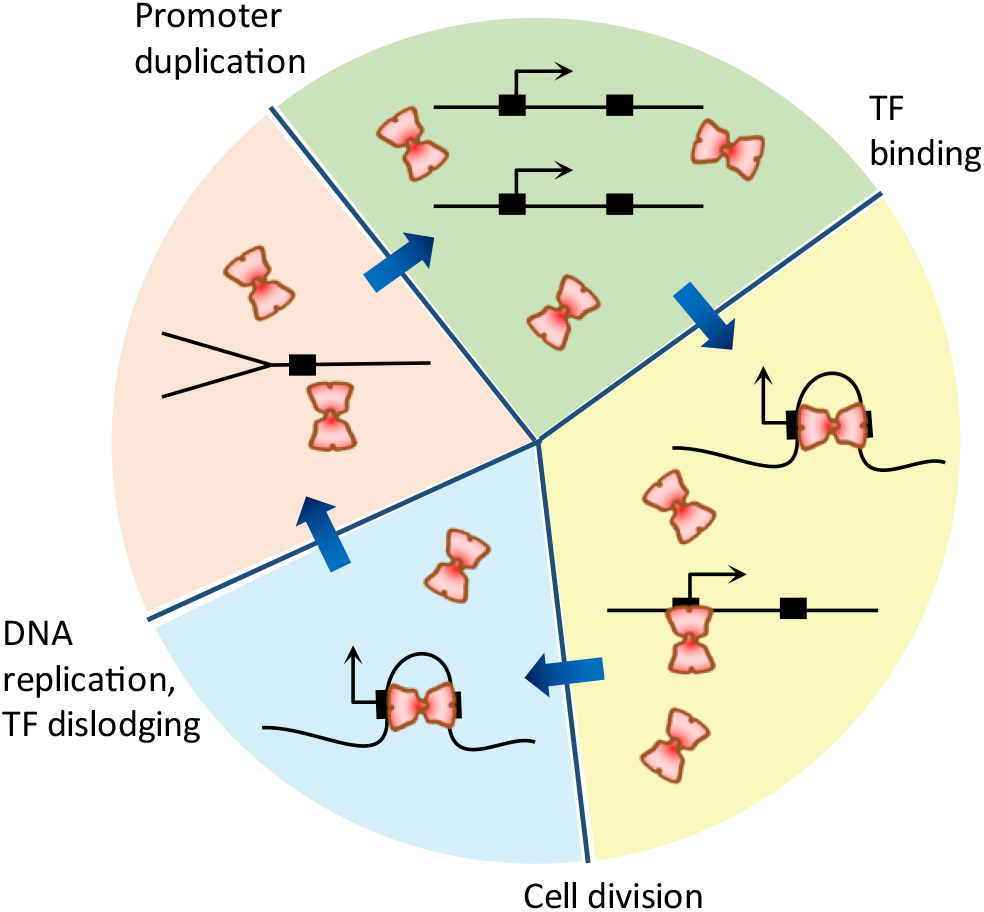
The cell cycle leads to strong recurrent nonequilibrium perturbations to gene regulation. The cartoon illustrates four sequential recurrent stages in gene regulation (clockwise direction). After cell division, there is a single copy of the promoter (black-angled arrow) and a set of TF binding sites (black rectangles) where the divalent TFs (red figures) can bind, potentially looping DNA (black line). DNA replication leads to dislodging of the TFs from the promoter region. After the passage of the replication fork, the promoter and DNA regulatory elements reappear in duplicate. Afterward, TFs should find their duplicated binding sites to reinstate regulation.

This paradoxical evidence is present prominently in the high efficiency observed in systems with just a few copies of the regulatory molecules, as in the *lac* operon with just ∼10 repressors per cell, and the plethora of experimental conditions which have been captured in detail with equilibrium models. The replication fork speed at about 1 kbp/s (Elshenawy et al., 2015; Pham et al., 2013) and size of 1.2kbp, as measured by the Okazaki fragment length (Balakrishnan and Bambara, 2013), indicate that in about 1 second the promoter and the regulatory elements are duplicated. Yet, the time for the repressors to find the operator site has been visualized to be about 30 seconds (Hammar et al., 2014), which would result in the production of ∼10 transcripts per promoter after replication at the usual transcription rate of 1 transcript every 3 seconds (Kennell and Riezman, 1977).

If the previous considerations were all true, repression would be 10 times less efficient than observed for the natural *lac* operon, which on average leads to fewer than 1 transcript per hour (Kennell and Riezman, 1977). Under these circumstances, equilibrium approaches would not work. Namely, without a much faster dynamics than observed, the kinetic limitations of the observed rates would prevent a very strong binding to work as effectively as needed before it is reset by the replication fork.

Therefore, the observed effectiveness of equilibrium approaches seems to underlie the experimentally observed efficiency. Strikingly, this paradoxical behavior is not just implied but has also been observed directly at the single-cell level, with transcription behaving as if the cell cycle were not present (Chang et al., 2022). These data showed not only that the completely repressed state in the natural, wild type (WT) system can be propagated across multiple generations without any transcriptional leakiness but also that the observed statistics of transcriptionally silent intervals and burst size statistics is consistent with the equilibrium dynamics quantified across independent experiments (Chang et al., 2022; Hammar et al., 2014).

Here, we develop a general theory for analyzing the impact of strong nonequilibrium perturbations on quasi-equilibrium systems and apply it to quantify the recurrent effects of cell cycle events in gene regulation systems. The results show that the cell cycle imposes strong limitations to the TF binding patterns that can be achieved over time and, at the same time, also provide an avenue, with short transient deactivation of transcription at the promoter after replication, that would allow the system to bypass these limitations. Interestingly, the *lac* operon is under both negative and positive control. Therefore, it is appealing to consider the potential role of CRP as the mechanism responsible for the delayed activation of the promoter. With values for the *in vivo* rates that are consistent across experiments, we show that the observed efficiency in synthetic systems that do not have active CRP falls at predictable nonequilibrium levels limited by the rate of dislodging of the TFs from DNA by the replication fork.

## Results

The development of the theory considers a state-based description of the system (Saiz and Vilar, 2006; Vilar and Saiz, 2013b) in which the fast-relaxing modes reach rapid equilibration between them (van Kampen, 2007) and collectively follow the slower dynamics of nonequilibrium states (Haken, 1983).

We consider the system characterized by an ensemble of *N* states, *S* = (0, 1, … *N* − 1). The reference state is indexed as *i* = 0, which we use also as the nonequilibrium state. The other states *i* > 0, with *i* ∈ *S*, are assumed to reach fast equilibration between them. The probability for the system to be in a state is denoted by *P*_*i*_ and the corresponding equilibrium value, by 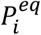.

### Quasi-equilibrium transcription

For the states *i* > 0 in quasi-equilibrium between them, the detailed balance principle (Beard and Qian, 2008) implies 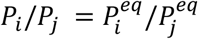, ∀*i,j* > 0, which together with the probability normalization expressed as Σ_*i*>0_ *P*_*i*_ = 1 − *P*_0_ leads to general expression

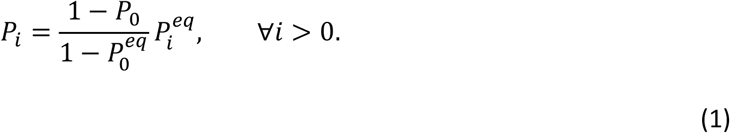

This expression is approach agnostic; namely, 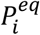 can generally be expressed in terms of equivalent TF and DNA properties, such as concentrations and equilibrium dissociation constants, concentrations and standard free energies, and number of molecules and free energy changes (Saiz, 2012).

The model of transcription considers that transcription at each state *i* takes place at a rate *γ*_*i*_ so that the overall transcription rate,

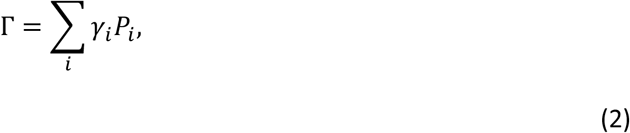

is expressed as the average over all the states. The quasi-equilibrium assumption for the states *i* > 0 implies 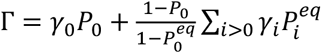, which can be rewritten as

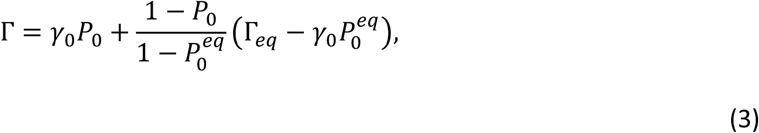

where 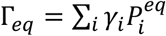 is the usual equilibrium transcription rate. This general expression we have obtained is extremely powerful as it provides an explicit quantification of the nonequilibrium transcription rate from its overall equilibrium value and the departure from equilibrium. Namely, the details of the intra quasi-equilibrium dynamics are not relevant provided that the overall effect is known.

### Nonequilibrium cell-cycle effects

We generally consider that the state *i* = 0 is out of equilibrium following the dynamics *dP*_0_/*dt* = −*k*_*in*_ *P*_0_ + *k*_*out*_ (1 − *P*_0_), indicating that the system can switch in and out of the state *i* = 0 with rates *k*_*in*_ and *k*_*out*_, respectively. In terms of a repressive system like the *lac* operon, this nonequilibrium state would be the promoter with no repressor bound to the operators, *k*_*in*_ would be the association rate of the repressor to any of the operators, and *k*_*out*_ would be the rate at which the operators end with no repressor. This equation can be rewritten as

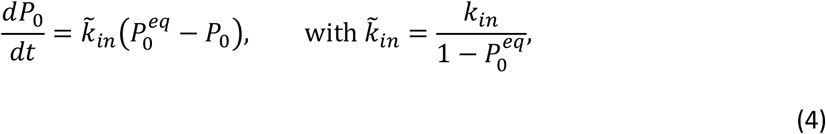

Noting that 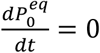 at equilibrium generally implies 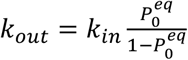.

To simplify the modeling of the cell cycle effects, we track just one of the daughter strands. We consider that the replication fork crosses the promoter at time 0 and that the effects are to clear the system from transcriptional regulators. In mathematical terms, this implies that at time zero *P*_0_ = 1 and *P*_*i*_ = 0, ∀*i* > 0, which results in

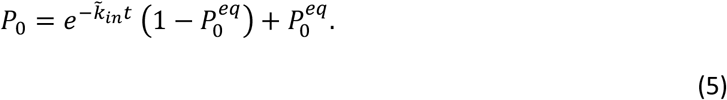

This dynamic characterization implies that the system will reach equilibrium on a time scale of 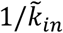. The dynamic evolution of the probabilities, *P*_*i*_, ∀*i* > 0, for the other states of the system follows straightforwardly from Eqs. (1) and (5).

To compute the average transcription 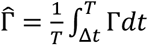 over a cell cycle of length *T*, we generally consider that transcription can resume after a time Δ*t* from the replication fork crossing the promoter, which after substitution of Eq. (5) in Eq. (3) leads to

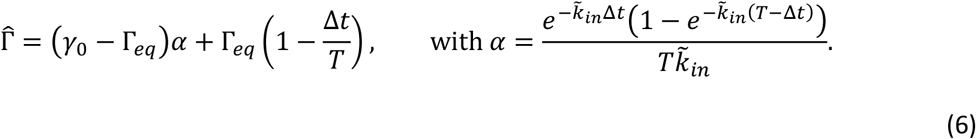

### Repression of transcription

The repression level, denoted by *R*, is the inverse normalized transcription and is formally defined as the maximal transcription, which corresponds to the fully unrepressed system when the repressor is not present or cannot bind to the promoter region, Γ_*max*_ = *γ*_0_(1 − Δ*t*/*T*), over the actual transcription 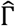:

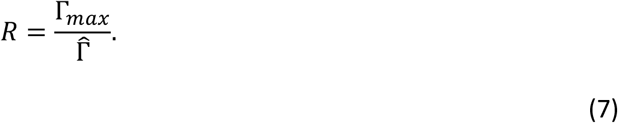

Combining the expression of the average transcription over a cell cycle with the expression of the equilibrium repression level *R*^*eq*^ = *γ*_0_/Γ_*eq*_ leads to the actual nonequilibrium repression level

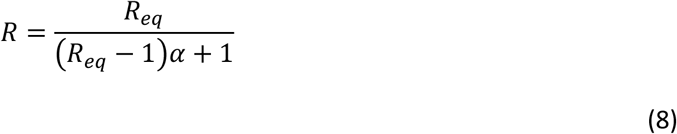

expressed in terms of the equilibrium value and the nonequilibrium parameter *α*.

In contrast to the equilibrium value, the kinetic analysis shows that the maximum achievable repression level is bounded from above by 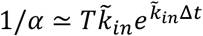 as a result of the cell-cycle nonequilibrium effects. In general, 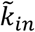 depends on the number of TF molecules, *n*_*TF*_, and under typical conditions, it is expected to increase proportionally to this number as 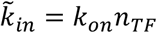, which makes the limit tunable at the cellular (e.g., regulation of the number of TFs), environmental (e.g. regulation of the binding of the TFs), and experimental (overexpressing or knocking down TFs) levels.

### Experimental validation with the lac operon

The *lac* operon has been the best and most comprehensively characterized genetic system since the discovery of gene regulation (Jacob and Monod, 1961). Its response to lactose relies on three regulatory DNA sites, a main operator O1 and two auxiliary operators O2 and O3, where the LacI repressor can bind. LacI’s binding to O1 prevents transcription by the RNA polymerase irrespective of its binding to O2 or O3 (Oehler et al., 1990). Allolactose, a derivative of lactose, and other analogous molecules prevent the specific binding of LacI, leading to the expression of the three genes involved in lactose metabolism. The auxiliary operators are required for the formation of a higher-order structure of DNA, a DNA loop between the main and an auxiliary operator bound simultaneously to a single LacI tetramer, to efficiently repress transcription using only a handful of LacI molecules. Looping is possible because LacI functions as a tetramer with two dimeric DNA binding domains (Lewis et al., 1996). As in the expression of other metabolic systems, the promoter needs activation by the complex CRP-cAMP to achieve significant transcription. The overall DNA looping-repressor-activator regulation motif is widely present among prokaryotes and eukaryotes (Cournac and Plumbridge, 2013).

To quantify the limitations imposed by the cell cycle on the naïve functioning of gene regulation systems, we apply the approach to the lac operon in *E. coli* with different combinations of operator deletions and sequence changes, WT and increased intracellular LacI numbers, changes in CRP activation, inhibition of repressors by inducers, and WT and synthetic promoters.

The scaling law, Eq. (8), connects equilibrium with actual values in terms of a single nonequilibrium parameter *α*. Therefore, we use previously computed equilibrium repression levels (Saiz and Vilar, 2008). For the value of *α*, we consider a characteristic value for the cell division time of *T* = 90 min and the association rate constant for the LacI tetramer binding to an operator of *k*_on_ = 0.3 molec^−1^min^−1^, as inferred from the bursting statistics of single-cell experiments at 30°C (Chang et al., 2022). The value of *k*_on_ is consistent with the upper and lower bounds established for the LacI dimer at 37°C and 25°C, respectively (Hammar et al., 2014) and is in line with those observed for other TFs in more complex organisms (Larson et al., 2011). Therefore, high repression, 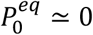, leads to 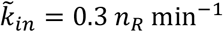, were *n*_*R*_ is the number of repressors.

### Cell-population activity assays

The early experiments that led to the discovery of gene regulation already measured that the activity of the *lac* operon increased more than 1,000-fold upon mutations in the DNA repressor region (Jacob and Monod, 1961). The key point is, in quantitative terms, that if transcription can resume as soon as the promoter is duplicated, the promoter would remain active, on average, for the value of 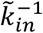; namely, 20 seconds for the WT case with 10 LacI tetramers per cell. Therefore, the TF kinetics limits the maximum repression level to 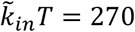 for WT, which is a factor 5 smaller than the observed value of 1,300 (Jacob and Monod, 1961; Oehler et al., 1990). In general, this limit would not significantly affect the single-operator synthetic systems with the auxiliary operators deleted, which, with a repression level of 20 (Oehler et al., 1990), is expected to be active for 4.5 min during the whole cell cycle. Nor would it have a substantial impact in the synthetic systems with unnaturally high numbers of LacI tetramers, such as those that express 900 LacI tetramers per cell with an estimated overall search time 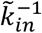 for the operators of 0.22 seconds.

The detailed analysis through Eq. (8) shows that, indeed, as the delay in resuming transcription increases, the behavior of the system becomes closer to the expected equilibrium value (Fig. 2). At WT number of repressors, the equilibrium value is generally higher than the measured one and there is a characteristic delay of about 40 seconds that exhibits the closest agreement with the measured values (Fig. 2). For systems with a single operator, repression is generally low and all the kinetic effects collapse within the equilibrium values for all range of repressor molecules per cell (Fig. 2).

**Figure 2.**
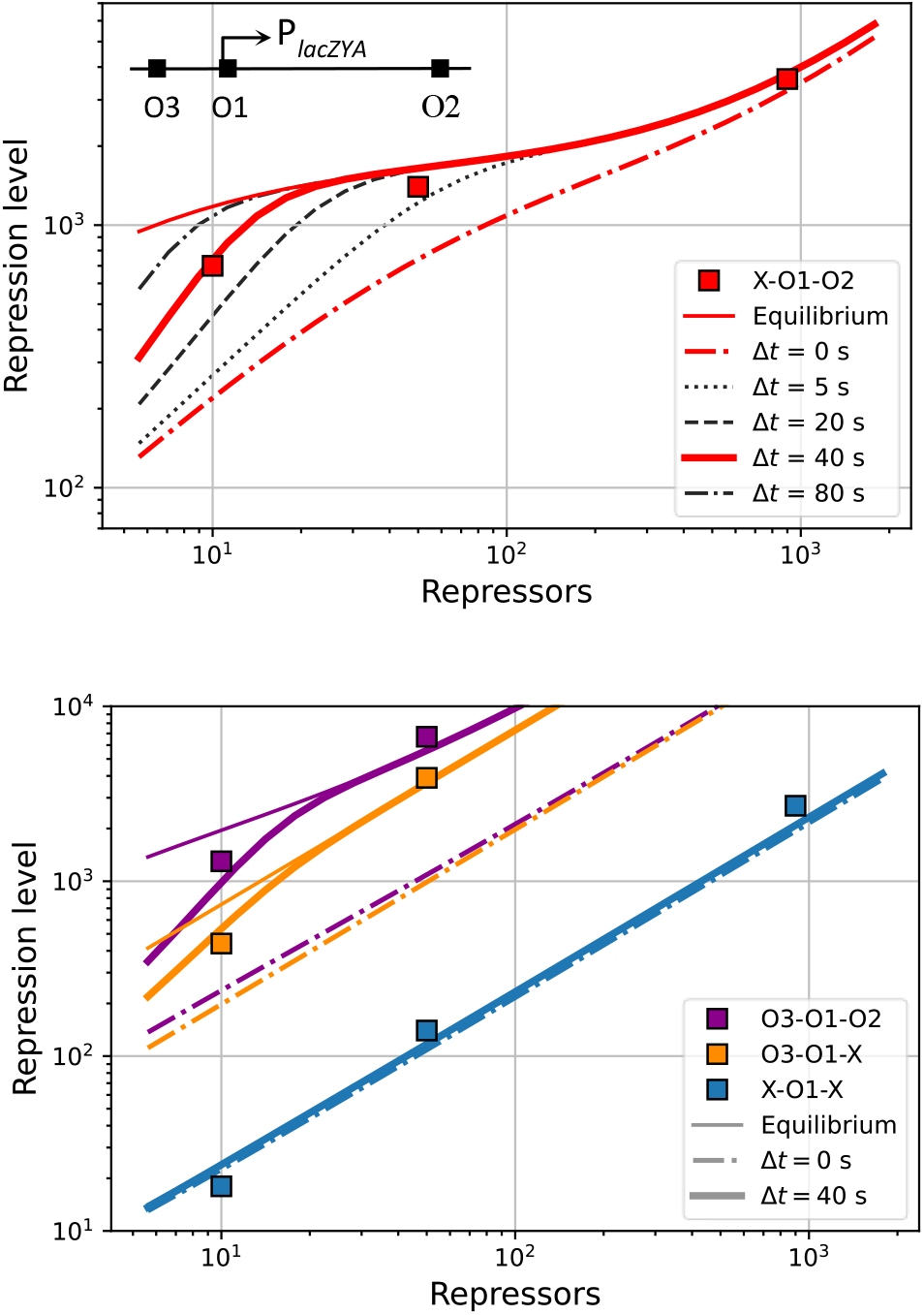
Transcription deactivation after replication for natural promoters makes regulation systems behave as if they were at equilibrium. Measured repression levels for the *lac* operon (Oehler et al., 1990) (colored square symbols) follow with minor differences the corresponding equilibrium predictions (Saiz and Vilar, 2008) (colored continuous thin lines) for different numbers of repressors per cell. The systems considered include the WT operator configuration (O3-O1-O2, bottom panel), O3 deletion (X-O1-O2, top panel), O2 deletion (O3-O1-X, bottom panel), and O2 and O3 deletions (X-O1-X, bottom panel). (The relative positions of the operators and promoter and shown on the top panel inset.) The nonequilibrium results from applying Eq. (8) to the equilibrium predictions with 40 seconds deactivation of transcription after replication (colored thick lines, Δ*t* = 40 *s*) are even closer to the measured repression levels. With no deactivation (colored dot-dashed lines, Δ*t* = 0 *s*), the nonequilibrium effects would prevent efficient repression. As the deactivation of transcription Δ*t* increases (multiple black lines, top panel), the system behaves as it were closer to equilibrium.

### *In vivo* single-transcript imaging

*In vivo* single-molecule imaging of transcript production from the *lac* system with its genes replaced by an RNA reporter has provided direct evidence of increased transcription after the replication fork (Wang et al., 2019). The probability of transcription in a 5-minute window after replication, with a value of 0.116, is about twice as much as that of previous and subsequent time windows, with an average value of 0.067. These values imply that 8.4% of the mRNA content has been produced upon replication. The observed values of ∼0.1 transcripts per cell with a degradation time for the transcripts of 2 min, results in a total mRNA production of 10 transcripts per cycle. Therefore, on average, one would expect the replication fork to have induced the production of Δ*m* = 0.84 transcripts. This quantity, with 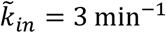 and a basal level of *γ*_0_ = 20 transcripts/min, leads to a delay in transcription reactivation of

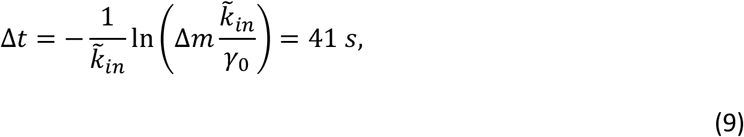

which closely follows those inferred in cell population experiments.

### Synthetic promoters without activator

The *lac* operon has both positive and negative control of transcription. Therefore, a potential source of delay in restarting transcription after replication is the activator CRP. We tested whether replacement of the natural CRP-dependent promoter for the *lac*UV5 promoter, which is constitutively active, revealed the underlying nonequilibrium effects.

The *lac*UV5 promoter has been widely used in multiple applications as it provides constitutively high transcription independently of CRP (Buc and McClure, 1985). Our analysis shows that if the system is far from the kinetic limit imposed by cell cycle perturbations, *R*_*eq*_*α* ≪ 1, both actual and equilibrium values are consistent with each other. Indeed, this is the case for the WT promoter with a delay in reactivating transcription of Δ*t* ≃ 40 s. If the delay is associated with CRP, discrepancies with the equilibrium values should be higher for the *lac*UV5 promoter, especially for regulation of the *lac*UV5 promoter with strong operators coupled through DNA looping at low to medium LacI numbers (Muller et al., 1996).

The single loop case for a main operator with a strong auxiliary operator leads to an equilibrium repression level,

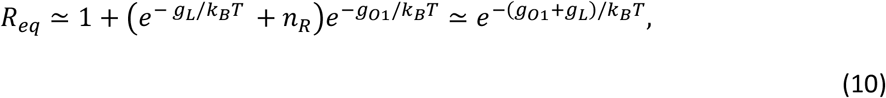

that is independent of LacI expression and the precise binding to the auxiliary operator (Vilar and Saiz, 2013b). It depends mainly on the free energy of binding to O1, *g*_*O*1_, and the free energy of looping, *g*_*L*_, which is substantially lower than −*k*_*B*_*T* In *n*_*R*_, with *k*_*B*_*T* being the thermal energy. However, this independence in LacI values is not observed for the experimental values of repression (Fig. 3).

**Figure 3.**
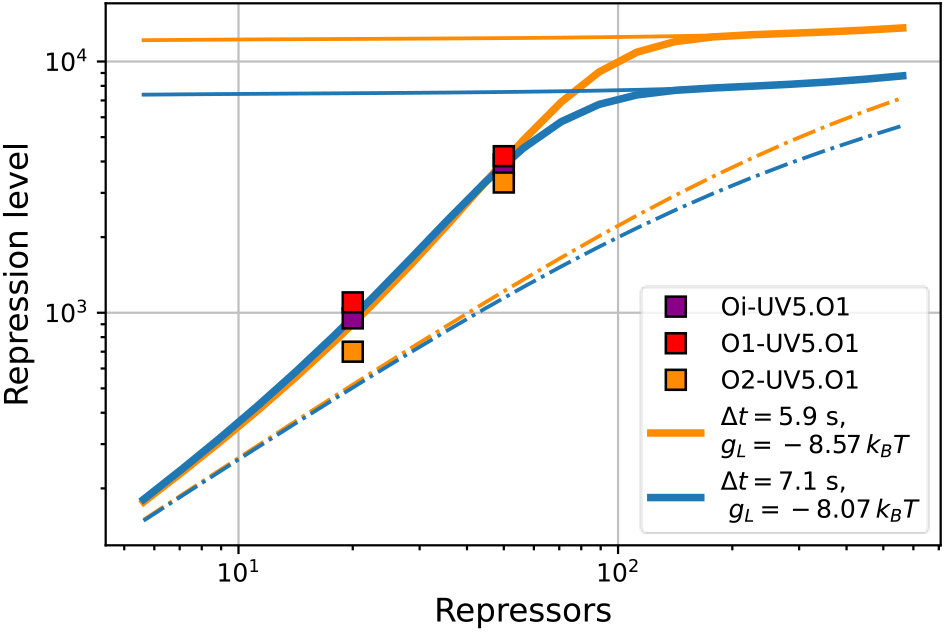
Regulation of the synthetic *lac*UV5 promoter is strongly determined by nonequilibrium effects. Measured repression levels (Muller et al., 1996) (colored square symbols) strongly diverge from the corresponding equilibrium predictions obtained through Eq. (10) (continuous thin lines: orange for *g*_*L*_ = −8.57 *k*_*B*_*T*, blue for *g*_*L*_ = −8.07 *k*_*B*_*T*) as the number of repressors per cell decreases. The systems considered include the *lac*UV5 promoter with O1 as the main operator for three different auxiliary operators [Oi (Oi-UV5.O1), O1 (O1-UV5.O1), and O2 (O2-UV5.O1)]. The nonequilibrium results from applying Eq. (8) to the equilibrium predictions with 5.9 seconds (*g*_*L*_ = −8.57 *k*_*B*_*T*) and 7.1 seconds (*g*_*L*_ = −8.07 *k*_*B*_*T*) deactivation of transcription after replication (thick lines) closely follow the measured repression levels. For low numbers of repressors, the repression levels converge to the results with no deactivation (dot-dashed lines, Δ*t* = 0 *s*).

The nonequilibrium repression level accurately reproduces the observed behavior and implies just a minor delay of 5.9 seconds in reactivating transcription for the WT values of *g*_*O*1_(= −0.85 *k*_*B*_*T*) and *g*_*L*_. The WT value of *g*_*L*_ = −8.57 *k*_*B*_*T* for the short O3-O1 loop should be expected to be a lower bound. It includes the stabilization effects of CRP-cAMP, which binds within the loop in the WT system but not in the synthetic *lac*UV5 promoter. The best agreement for the strongest auxiliary operator (Oi) replacing O3 is achieved with an increased free energy of looping of *g*_*L*_ = −8.07 *k*_*B*_*T* and a similar delay of 7.1 seconds. This increase in free energy is consistent with an upper bound for the missing stabilization of the loop by binding of CRP to its DNA site between O1 and O3 (Hudson and Fried, 1990). In both cases, the discrepancy between actual and equilibrium values of the repression level is about a factor 10 for low numbers of repressors (20 LacI tetramers), whereas it is negligible for the high number of repressors (50 LacI tetramers) after incorporating the lack of stabilization of the O3-O1 loop by CRP-cAMP.

### Induction without CRP-cAMP activation

The role of CRP in the *lac* operon has been a long-standing question. In the earlier literature, it was assumed that levels of the complex CRP-cAMP, the active form that binds DNA, were high in the absence of glucose and low otherwise (Pastan and Perlman, 1970), but subsequent studies found that the major mechanism from preventing the activation of the *lac* operon in the presence of glucose is inducer exclusion; namely, glucose inhibits the transport of lactose inside the cell (Inada et al., 1996). Consistently with these findings, it has been shown in precise detail that CRP-cAMP is naturally present at saturating levels under multiple growing conditions, including either glucose and glycerol, and that it enhances transcription by a factor ∼200 for fully induced systems and by a factor ∼50 for fully repressed systems (Kuhlman et al., 2007).

To analyze the induction process in synthetic systems without cAMP, we use the repression values of the system with saturating exogenous cAMP levels (Kuhlman et al., 2007), which closely mimics the WT case, as a first approximation to the equilibrium values *R*_*eq*_. Binding of one IPTG inducer molecule to one LacI monomer deactivates the specific DNA binding of the corresponding dimeric domain (Oehler et al., 2006). Considering *p*_*F*_ = 1/(1 + [*IPTG*]/*K*_*I*_) as the probability that one monomer is free from inducer, the number of fully active LacI tetramers is 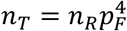 and the number of tetramers with one binding domain impaired is 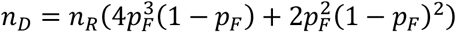, which leads to

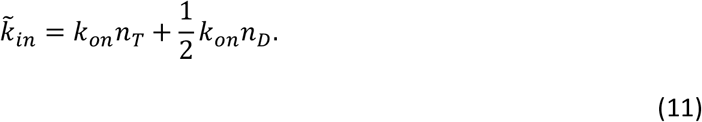

As value of the inducer dissociation constant, we use explicitly *K*_*I*_ = 7.2 μM from the experimentally determined tight range of 6.4 – 8.2 μM (Oehler et al., 2006), which closely reproduces multiple equilibrium and kinetic induction situations (Vilar and Saiz, 2013a). The resulting active transcription after replication (Δ*t* = 0) show an almost perfect agreement with the observed repression level (Fig. 4).

**Figure 4.**
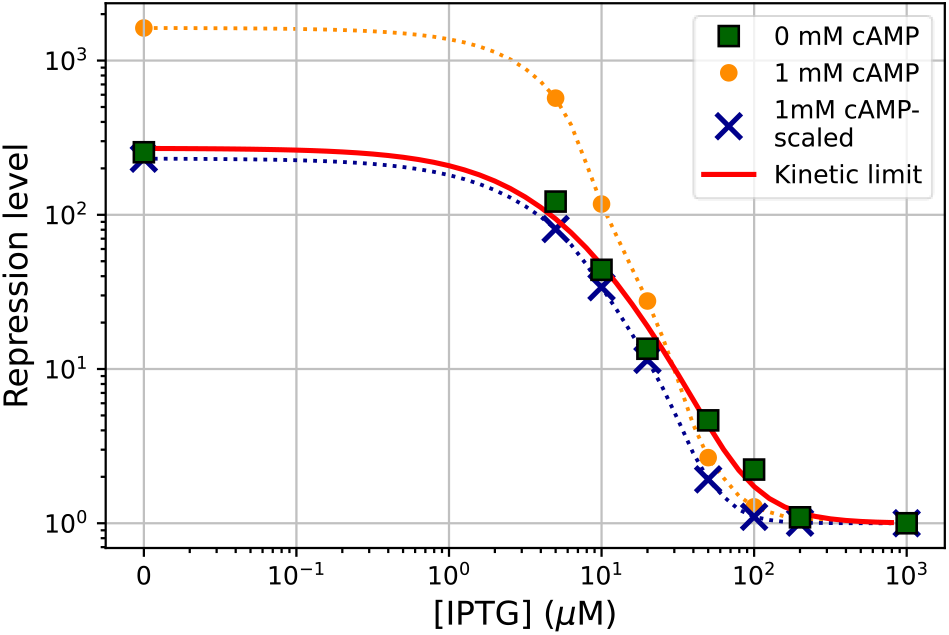
Regulation of the natural *lac* promoter without CRP activation is kinetically determined by nonequilibrium effects. The measured repression levels for strain TK310 with 0 mM cAMP (Kuhlman et al., 2007) (square symbols) for different inducer concentrations are closely approximated by scaling the measured repression levels with 1 mM cAMP (Kuhlman et al., 2007) (yellow dots), as proxies of *R*^*eq*^, through Eq. (8) with Δ*t* = 0 (crosses). The kinetic limit (red continuous line), obtained with *R*^*eq*^ = ∞ and Δ*t* = 0 in Eq. (8), closely follows the measured repression for TK310 with 0 mM cAMP.

Because the observed repression values for the system with saturating values of cAMP are expected to be lower than the corresponding equilibrium values, we computed the kinetic limit of the repression level as well (Fig. 4). Strikingly, the kinetic limit provides a perfect match to the experimental values, indicating that the binding at the promoter is sufficiently long-lived for the induction process to be entirely determined by the recurrent nonequilibrium kinetics induced by dislodging of the repressor by the DNA replication fork.

### Quantification of the transcription reactivation delay

Overall, our analysis has shown that promoters positively regulated through CRP-cAMP behave as if they were at equilibrium whereas promoters that lack this activation behave strongly limited by the nonequilibrium dynamics resulting from the replication fork dislodging the associated TFs. Precise quantification of different *lac* promoter and operator configurations in *E. coli* (Fig. 5) show that the equilibrium-like behavior is associated with transcription being inactive after replication for 39.7 seconds on average and that nonequilibrium effects are prominent for fast restoring of transcription after replication, with an average delay of 4.4 seconds. These values are higher and lower, respectively, than the 20 seconds 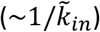 needed by the set of 10 TFs to find a functional target on average.

**Figure 5.**
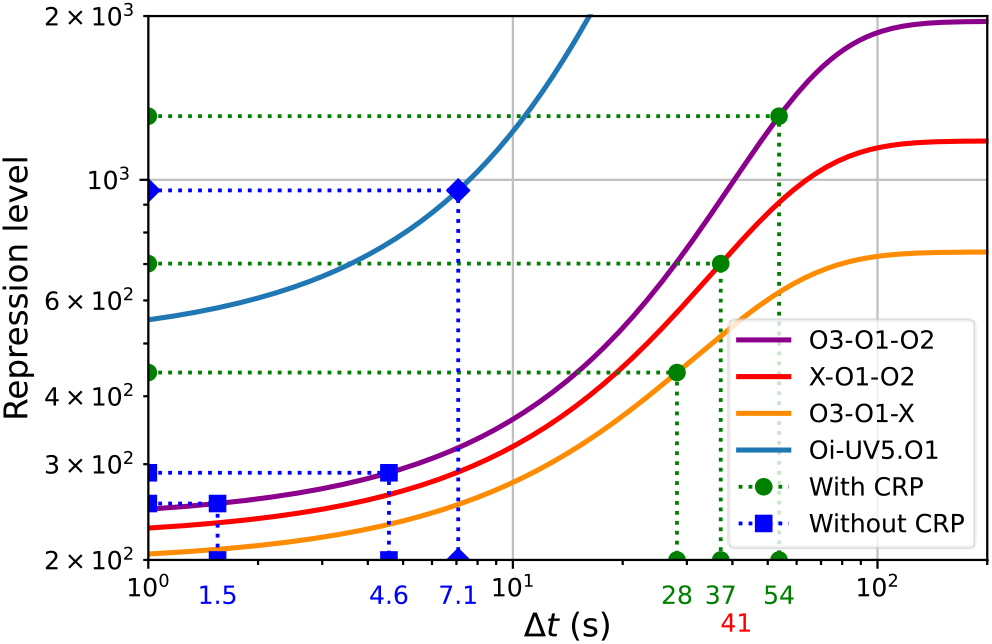
Transcription deactivation after replication is 10-fold longer in WT regulation than in the absence of active CRP. The duration of transcription deactivation is obtained through Eq. (8) as the time Δ*t* at which the computed repression level (continuous colored lines) coincides with the measured value (symbols). O3-O1-O2, X-O1-O2, and O3-O1-X correspond to systems in Fig. 2 with 10 repressors/cell (green dots); Oi-UV5.O1 corresponds to the system in Fig. 3 with 20 repressors/cell (blue diamonds). O3-O1-O2 systems without active CRP (blue squares) include strains TK230 and TK310 (0 mM cAMP) (Kuhlman et al., 2007). Deactivation times Δ*t* for WT regulation (green values; ⟨Δ*t*⟩ = 39.7 *s*) are on average 10-fold longer than for regulation without active CRP (blue values; ⟨Δ*t*⟩ = 4.4 *s*). The deactivation time inferred from single-transcript imaging (red value) agrees with the values for WT regulation.

## Discussion

Naturally occurring genetic control systems reliably function through the recurrent perturbations of the cell cycle, including nonequilibrium phenomena as diverse as dislodging of TFs from DNA by the replication fork, duplication of the gene, and partition of the TFs into daughter cells. The *lac* operon in *E. coli*, a paradigm of efficiency in gene regulation and the best characterized genetic system, can do so with just a few repressor molecules per cell.

The general theory we have developed has provided a general avenue to analyze the effects of nonequilibrium perturbations in gene regulation. Applied to the cell cycle effects, our results have led to an explicit scaling law that relates observed transcription with its expected equilibrium value in terms of a single parameter that combines the slowest time scale of the system and the generation time. This expression has revealed the existence of major physical constraints in the limits of repression, showing that efficient repression is severely limited by the kinetics of the TFs. Paradoxically, mechanistic predictive models accurately account for a substantial body of the observed average-cell behavior, but these models remarkably fail when they are improved to consider the molecular perturbations associated with the cell cycle (Chang et al., 2022). In this regard, the *lac* operon is operating 10 times more efficiently than naïvely allowed by the mechanisms considered up to now.

We have identified a class of situations in which it is possible to reconcile equilibrium models with the observed behavior by considering that transcription gets momentarily deactivated upon DNA replication. In the *lac* operon, these situations include the natural system and other situations with the WT *lac* promoter under the positive control of CRP-cAMP. In contrast, regulation without an adjuvant activator, as we have shown for both the *lac*UV5 promoter and systems without CRP-cAMP activation, is generally associated with results that are not compatible with equilibrium approaches, up to a factor 10 as observed experimentally, and that need to consider nonequilibrium cell cycle effects. In general, inefficient systems are not subject to these limitations, as for instance, artificial promoters regulated with weak sites. In these cases, the effects of the cell cycle are expected to be present but get masked by the relatively high overall transcription at low numbers of TFs and are not present at high numbers of TFs, which lead to a fast dynamics resulting from high concentrations.

We have analyzed two lines of evidence, cell-population activity assays and *in vivo* transcript imaging, in the light of our approach to quantify the deactivation of transcription conclusion. These two completely different approaches provide the same results.

If the TFs had a much faster kinetics, the nonequilibrium effects would not be apparent. The experimental evidence, however, shows that they do not. The main proof is the direct measurement of the *in vivo* association and dissociation rate constants for the LacI repressor dimer (Elf et al., 2007; Hammar et al., 2014). Concomitantly, the time scale of repressor search kinetics has been confirmed by the statistics of transcriptionally silent intervals and protein burst sizes in single cells for multiple regulatory setups (Chang et al., 2022).

Further supporting the main conclusion, we have shown that highly repressible regulatory systems without CRP-cAMP activation are not consistent with the expected equilibrium behavior, which would rule out much faster kinetics (Figs. 3 and 4). In addition, in these systems, the results get closer to the equilibrium values when the number of LacI molecules increases, as expected as the overall TF target search times decrease (Fig. 4).

Deactivation of transcription after replication has fundamental implications at the physiological level. It has been observed in living cells at the single-molecule level that infrequent events of complete dissociation of LacI from DNA result in large bursts of permease expression that trigger induction of the *lac* operon and that partial dissociation of LacI from looped DNA leads only to basal expression (Choi et al., 2008). Therefore, the *lac* operon without deactivation of transcription would be induced frequently after replication rather than stochastically and infrequently. In more complex organisms, like yeast and other eukaryotes, it is well established that there are long delays in restarting transcription after DNA replication and that the delay depends on the type of gene (Stewart-Morgan et al., 2019). Therefore, the suppression of the effects of major nonequilibrium perturbations through transcriptional deactivation could also be present in these more complex systems.

Overall, our results show that the observed consistency with equilibrium approaches is not because of a fast dynamic behavior that moves the system close to equilibrium sooner but the consequence of the underlying cellular strategy of shutting down transcription after replication for as long as nonequilibrium effects are present. In this regard, the observed unreasonable effectiveness of equilibrium approaches in natural systems is not coincidental but the result of an efficient, beneficial behavior selected through evolution, as other key properties of natural systems (Dekel and Alon, 2005).

## Acknowledgments

J.M.G.V. acknowledges support from Ministerio de Ciencia e Innovación under grants PGC2018-101282-B-I00 and PID2021-128850NB-I00 (MCI/AEI/FEDER, UE). L.S. is indebted to the Max Planck Institute for the Physics of Complex Systems for hosting her for a sabbatical stay and to the ELBE visiting faculty program.

## Author contributions

J.M.G.V and L.S conceived, designed, and performed the research.

